# Predators affect a plant virus through direct and trait-mediated indirect effects on vectors

**DOI:** 10.1101/2021.02.17.431666

**Authors:** Benjamin W. Lee, Robert E. Clark, Saumik Basu, David W. Crowder

## Abstract

Arthropods that vector plant pathogens often interact with predators within food webs. Predators affect vectors by eating them (consumptive effects) and by inducing antipredator behaviors (non-consumptive effects), and these interactions may affect transmission of vector-borne pathogens. However, it has proven difficult to experimentally tease apart the effects of predators on vector fitness and behavior as they are often correlated. We addressed this problem by assessing how both aphids and an aphid-borne pathogen were affected by variable predation risk. Specifically, we experimentally manipulated ladybeetle predators’ mouthparts to isolate consumptive, and non-consumptive, effects of predators on aphid fitness, movement, and virus transmission. We show that although lethal predators decreased aphid vector abundance, they increased pathogen transmission by increasing aphid movement among hosts. Moreover, aphids responded to risk of predation by moving to younger plant tissue that was more susceptible to the pathogen. Aphids also responded to predator risk through compensatory reproduction, which offset direct consumptive effects. Our results support predictions of disease models showing alterations of vector movement due to predators can have greater effects on transmission of pathogens than vector consumption. Broadly, our study shows isolating direct and indirect predation effects can reveal novel pathways by which predators affect vector-borne pathogens.

## Intro

Within food webs, arthropod vectors that transmit plant pathogens engage in direct and trait-mediated indirect interactions with individuals of other species such as competitors, mutualists, and predators (Clark et al. 2019, Crowder et al. 2019). Models and empirical studies show that species such as predators and competitors affect the spread of vector-borne pathogens through direct and trait-mediated indirect effects on vectors (Finke, 2012, Chisholm et al. 2019, Clark et al. 2019). For example, predators may interfere with pathogen transmission by killing vectors (consumptive effects) but may increase pathogen spread if vectors increase their movement when predators are present (non-consumptive effects) (Crowder et al. 2019). However, experimentally untangling the direct and indirect effects of predators on vectors and vector-borne pathogens has proven difficult, given that consumptive and non-consumptive effects are not independent.

Most insect vector species are attacked by predators; if predators only killed vectors, they should decrease vector-borne pathogen transmission by reducing vector abundance (Moore et al. 2010). However, in response to predators, insect vectors exhibit a range of behaviors such as dropping from plants (Losey and Denno, 1998, Fan et al. 2017). Predation risk can also affect vector feeding behavior and movement (Smyrnioudis *et al*., 2001, Hodge *et al*., 2011, Kersch-Becker & Thaler, 2015, Tholt et al. 2018). While such behaviors are effective ways to defend again predation, they may also come at a cost of reduced feeding duration and diet quality, and lower reproductive output for vectors (Preisser *et al*., 2007, Jones and Dornhaus, 2011). However, while reviews show that the non-consumptive effects of predators are often equally or more important in affecting prey demographics as consumptive effects (Preisser *et al*. 2005), few studies have isolated how direct (consumptive effects) and trait-mediated indirect (non-consumptive effects) of predators might affect vector-borne pathogens.

Direct predator effects on vector-borne pathogens are expected to be straightforward, with reduced vector abundance slowing pathogen transmission. However, indirect effects of predators on these same transmission mechanisms are more complex. Models suggest that increased vector movement between host plants due to predation risk should accelerate virus transmission, while reduced vector feeding duration should slow transmission (Finke, 2012, Crowder *et al*. 2019). Empirical support for this has been shown in a system where predators increased transmission of *Barley yellow dwarf virus* in wheat by increasing movement of the aphid vector (Smyrnioudis et al. 2001). In contrast, when predators interrupted feeding by aphid vectors without affecting host-to-host movement, virus transmission was slowed (Long & Finke 2015). While informative, these studies had variation in vector abundance, and did not measure how vector abundance and behavior interactively affected transmission. This highlights the difficulty in isolating tradeoffs between consumptive and non-consumptive predator effects on pathogens, and a need for more targeted assessments of how vector responses to predation risk affect pathogen transmission.

In this study, we addressed these knowledge gaps in a model plant pathosystem comprised of the aphid vector *Acyrthosiphon pisum*, the host *Pisum sativum*, the pathogen *Pea-enation Mosaic Virus* (PEMV), and the ladybeetle predator *Hippodamia convergens*. We experimentally isolated consumptive (direct) and non-consumptive (indirect) predator effects by manipulating both predator presence and predator lethality. In response to these treatments, we measured how aphid vector populations, and their capacity to transmit the PEMV pathogen, responded to variable lethal and non-lethal predation risk. Data from our experiments were incorporated into structural equation models to isolate the direct and trait-mediated indirect pathways by which predators affected aphid abundance, aphid movement, and pathogen transmission.

## Material and Methods

### Insect and Virus maintenance

Pea aphid (*Acyrthosiphon pisum* Harris) colonies were originally collected in commercial pea fields in Washington State, and were maintained on potted pea plants (*Pisum sativum* L. cv. “Banner”) in greenhouses in Pullman, WA, USA under the following environmental conditions: 23 ± 2°C, L16:D8 photoperiod. Our PEMV isolate was obtained from the University of Idaho and was maintained by transferring pea aphids fed on PEMV-infected pea plants into uninfected pea aphid colonies, introducing clean plants as needed. Samples from infectious and uninfectious pea aphid colonies were tested monthly for the presence of PEMV using RT-PCR; these samples confirmed nearly 100% infection levels in the infectious colony and 0% in the uninfectious colony. *H. convergens* predators were collected from pea and alfalfa fields in Washington State 7 days prior to experiment start and stored in a growth chamber at 4 °C until needed.

### Effects of Predation Risk on Vectors and PEMV Prevalence

To structure our examination of predation effects on pea aphids (both adults and nymphs) and PEMV, we developed an *a priori* interaction network (Fig. 1). We predicted both lethal and risk predator treatments would reduce aphid abundance and increase movement. While reduced aphid abundance should decrease PEMV prevalence, increased aphid movement may increase PEMV prevalence by causing vectors to contact more hosts (Finke, 2012, Crowder et al. 2019). We expected vector movement to affect PEMV prevalence more than abundance (Chisholm et al. 2019), resulting in a net indirect positive effect of predators on PEMV prevalence.

**Figure 1.**
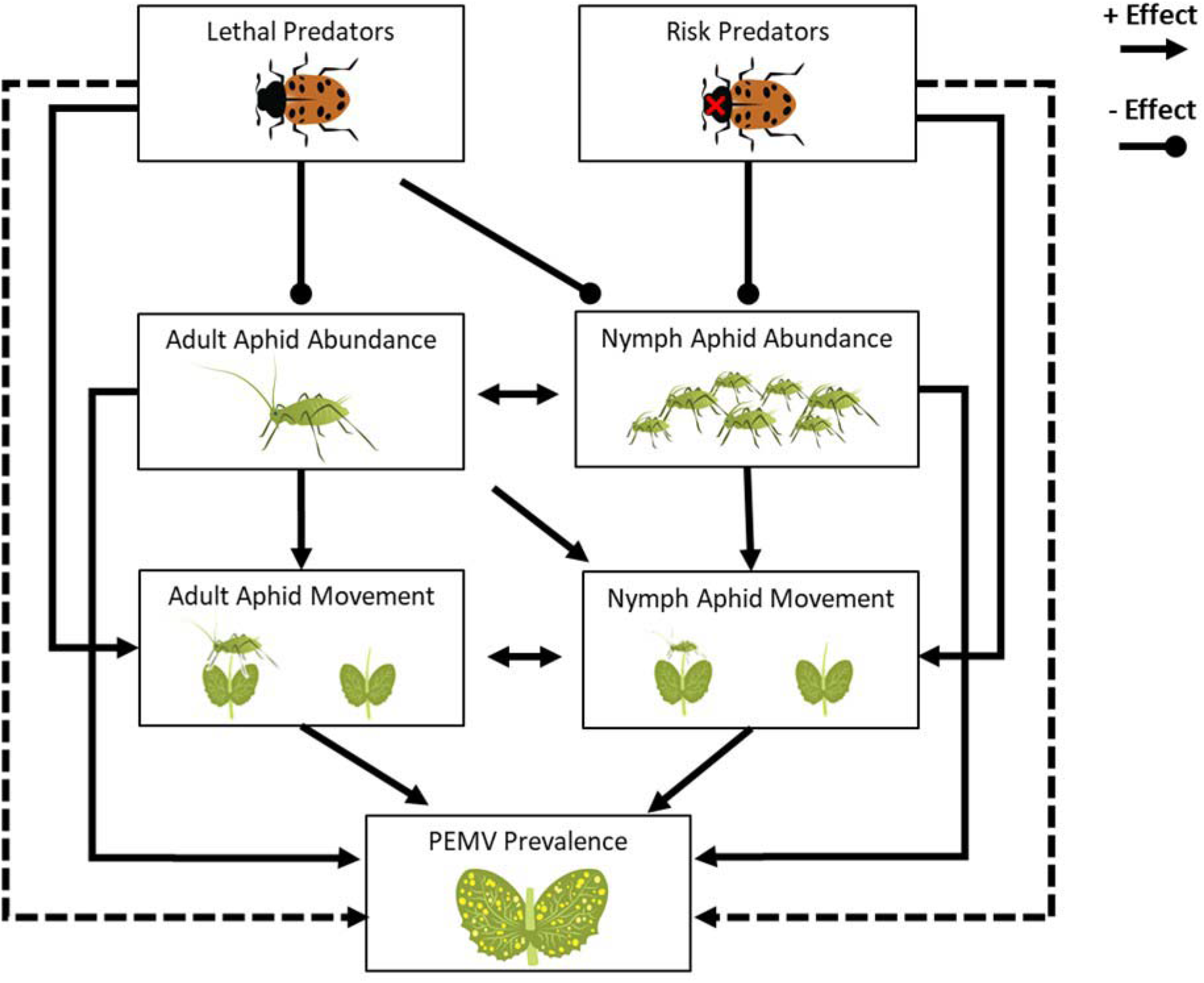
Interaction network with *a priori* predictions about indirect effects and the direction of effects (positive/negative). Boxes show predator treatments and response variables (adult/nymph aphid abundance/distance from center plant, and PEMV prevalence). We predicted that predation risk would directly reduce aphid abundance and increase aphid movement (full lines), indirectly increasing PEMV prevalence (dotted lines).

To test predictions of our *a priori* model, we conducted a field mesocosm experiment. By manipulating predator mouthparts, we were able to isolate the consumptive and non-consumptive predator effects on the movement and reproduction of vectors, and PEMV transmission. The experiment was conducted on bare-soil plots at the Palouse Conservation Farm in Pullman, WA, USA in two blocks (June, July 2018). Pea plants (*P. sativum* L. cv. “Banner”) were planted in 10 cm pots in the greenhouse in potting soil (Sun Gro® Sunshine® LC1 Grower Mix) 2 wk before the experiment. For each replicate, 25 potted plants were buried in bare soil in a 5 × 5 square grid within 2 × 2 × 2m cages with amber mesh Lumite screening (Bioquip, Gardena, California, USA), with 20cm of space between the centers of each pot. Cages were buried 3in below the soil surface to prevent escape of organisms, and peas were completely buried inside their pots to provide an even, contiguous surface throughout the mesocosm, reflecting field conditions.

Each replicate was randomly assigned one of three treatments: Control (no predators); Risk (4 non-consumptive “risk” *H. convergens*); or Lethal (4 unmanipulated “lethal” *H. convergens*). This density reflects ladybeetles observed in commercial pea fields (Lee, personal observation). Lady beetles were held individually at 25°C in petri dishes and fed *A. pisum* ad-libitum for 72 h, then starved for 48 h before receiving treatments to standardize hunger levels. “Risk” predators were briefly anesthetized with CO_2_ and a small droplet of clear nail polish was applied to seal their mandibles, with care taken not to restrict palps or antennae. This treatment prevents lady beetles from killing and consuming aphids but allows for full movement and prey-seeking behavior (Kersch-Becker & Thaler, 2015). “Lethal” predators were also anesthetized and received a droplet of polish on the elytra. Predators were then stored at 4°C overnight before use.

In each cage, 25 7-d old adult PEMV-infectious pea aphids were confined within a fine-mesh frame on the centermost pea plant for 24 h. After 24 h, the mesh was removed, established pea aphids were counted, and predators were released into the mesocosm. Adult and nymph pea aphid populations on each plant were recorded after 5 d. Missing or dead predators were replaced daily. After 5 d, aphids were removed with an aspirator and plants were treated with a granular formulation of imidacloprid (Bayer Crop Science, NJ, USA) to kill remaining aphids and cease virus transmission. Plants then grew for 5 d to accumulate viral titer before being visually evaluated for PEMV symptoms. For each replicate, aboveground tissue from 3 pea plants from each mesocosm quadrant were destructively sampled (Fig. S1), frozen in liquid nitrogen, and stored at −80°C. PEMV titer from the 3 plants in each quadrant was determined using rtPCR and quantified with ImageJ (US NIH, Bethesda, Maryland, USA) to measure electrophoresis gel band intensity relative to a positive control (Fig. S2). These pooled titer measurements were used to validate visual evaluations of PEMV presence within each mesocosm, as quantifying titer for all individual plants was unfeasible. Average PEMV titers, however, were highly correlated with visual evaluations of PEMV prevalence (Pearson’s Correlation, *r*_*46*_ = 0.80, *P* < 0.001). Eight replicates for each treatment (Control, Risk, Lethal) were conducted per block.

### Effects of Predation Risk on Vector Feeding Location

We conducted a second greenhouse experiment to further assess risk and lethal predator effects on vectors, focusing on individual aphid feeding location. Pea plants were grown for 2 weeks, and pea aphids were raised to adulthood on PEMV-infected peas as previously described. 5 pea plants were arranged in a row within 0.3 × 0.3 × 0.6m black mesh cages (Bioquip, Gardena, California, USA) and potting soil was spread across the tops of pots to provide a contiguous surface. ‘Risk’ and ‘Lethal’ *H. convergens* predators were also prepared as described, starved for 48h, and held at 4°C for 24h before use. 15 PEMV-infectious pea adult aphids were confined on the center plant in each row by a mesh bag for 24 hours before the bag was carefully removed and 3 ‘Risk’ or ‘Lethal’ *H. convergens* predators were added. The number, host plant, and feeding locations (top or bottom half of plant) of aphid pea adults and nymphs was recorded daily for three days. Sixteen replicates were conducted for each treatment.

### Data Analyses

We used structural equation modeling (SEM) to test *a priori* predictions for the field mesocosm experiment, using predator treatment (‘Risk’ or ‘Lethal’) and block as predictors and pea aphid abundance, pea aphid movement, and PEMV prevalence as responses (Fig. 1). Abundance was the total pea aphid adults or nymphs in the entire mesocosm, and movement was the average distance of aphid adults or nymphs from the center release plant. Parameters were continuous (aphid abundance and movement) or binary counts (number of plants infected out of 25). Block was included in all models as a fixed effect, as hotter temperatures in the second block reduced aphid abundance (GLM, *t*_*47*_ = −5.13, *P* < 0.001). In our analysis, non-significant paths were dropped if doing so reduced AIC, and paths were added if models without them were rejected by tests of direct separation (Lefcheck 2016). Predictor coefficients were standardized by their standard deviation to allow for comparisons of effects on different responses (Fig. 2, Table S1).

**Figure 2.**
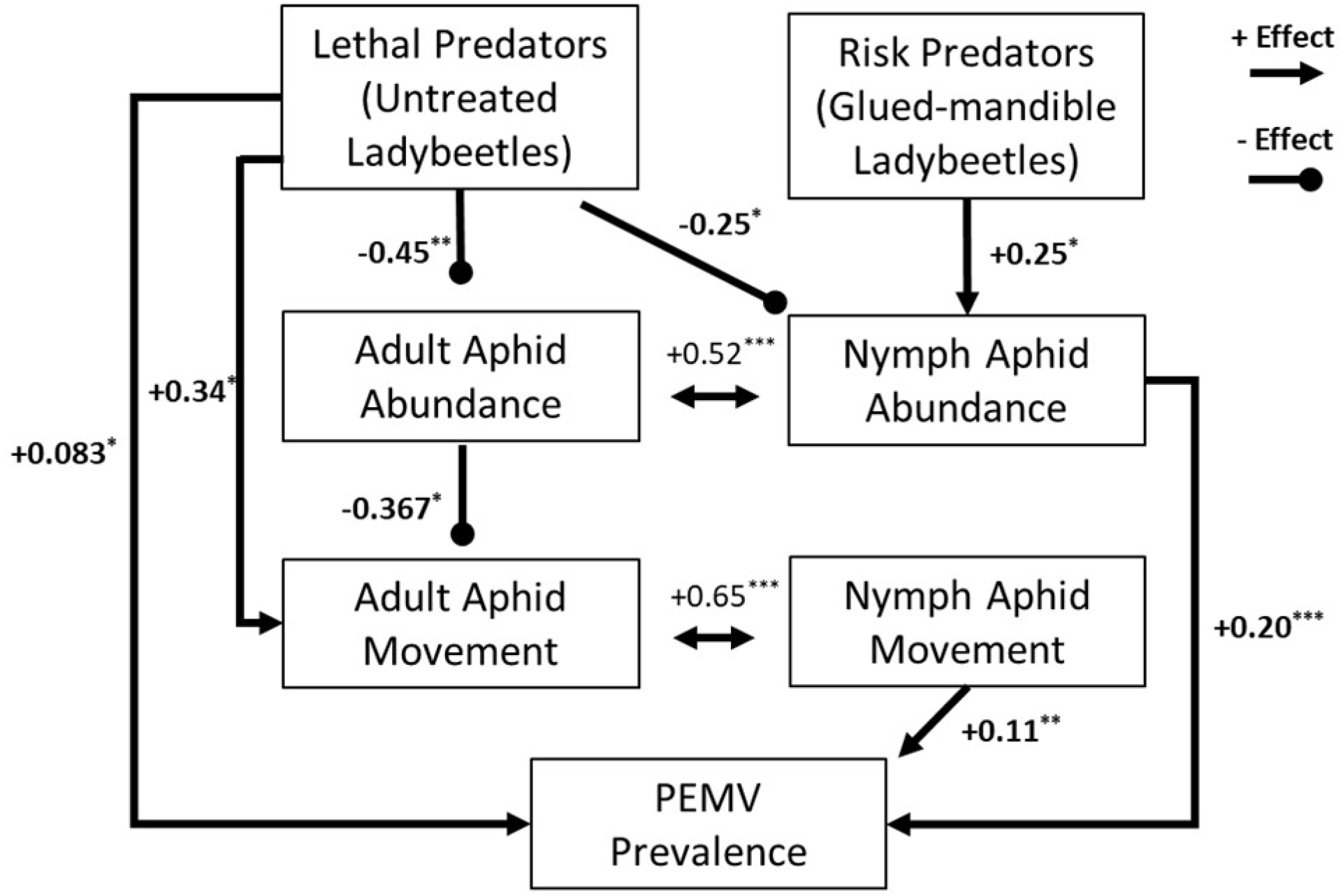
Accepted model from confirmatory path analysis (P = .87, Fischer’s C = 13.154, df = 20). Bidirectional arrows indicate correlated errors. Lethal predators increase PEMV prevalence directly (P = .02, Table S1) while decreasing PEMV through density-mediated indirect interaction.

We ran a series of generalized linear mixed models (GLMMs) to evaluate aphid responses to predators in the greenhouse experiment, using predator treatment and day as fixed effects, cage as a random effect, and adult and nymphal aphid abundance, distance from center plant, and proportion feeding on top vs. bottom half of host plant as responses. The fit of GLMMs to the observed data distributions were controlled by inspecting residuals and QQ plots. Binomial models for feeding location were weighted by total aphid abundance in each mesocosm to account for differences in abundance between treatments. All data analyses were conducted using R v 3.5.2 (R Working Group, 2018), using the packages ‘lme4’ for GLMMs (Bates *et al*. 2015), and ‘piecewiseSEM’ for structural equation models (Lefcheck, 2016). Posthoc analyses were conducted using the ‘emmeans’ package (Lenth 2015) and significance tests were based on analysis of deviance χ^2^ tests using the ‘car’ package (Fox and Weisberg 2019).

## Results

### Effects of Predation Risk on Vectors and PEMV Prevalence

Plants were 25.4% and 35.9% more likely to be infected with PEMV than controls in lethal and risk predator groups, respectively (χ^2^ = 4.93, df = 2, *P* = 0.085), and our best-fit SEM (*P* = 0.87, Fishers’s *C* = 13.2, df = 20) showed direct and indirect pathways by which predators affected aphids and PEMV (Fig. 2). Lethal predators reduced adult (βstd = −0.45, df = 46, *P* = 0.001) and nymph abundance (βstd = −0.25, df = 44, *P* = 0.037) but increased adult aphid movement (βstd = 0.34, df = 44, *P* = 0.050). Contrary to predictions, risk predators increased nymph abundance (βstd = 0.25, df = 44, *P* = 0.041). Across aphid responses, adult aphid movement decreased when adult abundance increased (βstd = −0.37, df = 43, *P* = 0.027), and there were positive correlations between adult and nymph abundance (βstd = 0.51, df = 48, *P* < 0.001) and adult and nymph movement (βstd = 0.65, df = 48, *P* < 0.001). PEMV prevalence was primarily driven by nymph abundance (βstd = 0.20, df = 43, *P* < 0.001) and nymph movement (βstd = 0.11, df = 48, *P* = 0.002), with a direct effect of lethal predator treatment (βstd = 0.08, df = 48, *P* = 0.023) (Fig. 2).

### Effects of Predation Risk on Vector Feeding Location

Predator treatments increased the proportion of aphid adults feeding on the top half of plants (χ^2^= 8.18, df = 2, *P* = 0.017), though this effect was less pronounced for nymphs (χ^2^ = 5.09, df = 2, *P* = 0.078) (Fig. 3, Table S2). Lethal predators reduced adult aphid abundance (t_136_ = −2.93, P = 0.0039) and nymphs over time (t_93_ = −7.26, *P* < 0.001) (Fig. S3a,b), and increased the dispersal of pea aphid adults (t_138_ = 3.01, *P* = 0.013) and nymphs (t_45_ = 5.10, *P* < 0.001) (Fig. S3c,d). Risk predators alone did not significantly affect aphid abundance or dispersal (Fig. S3).

**Figure 3.**
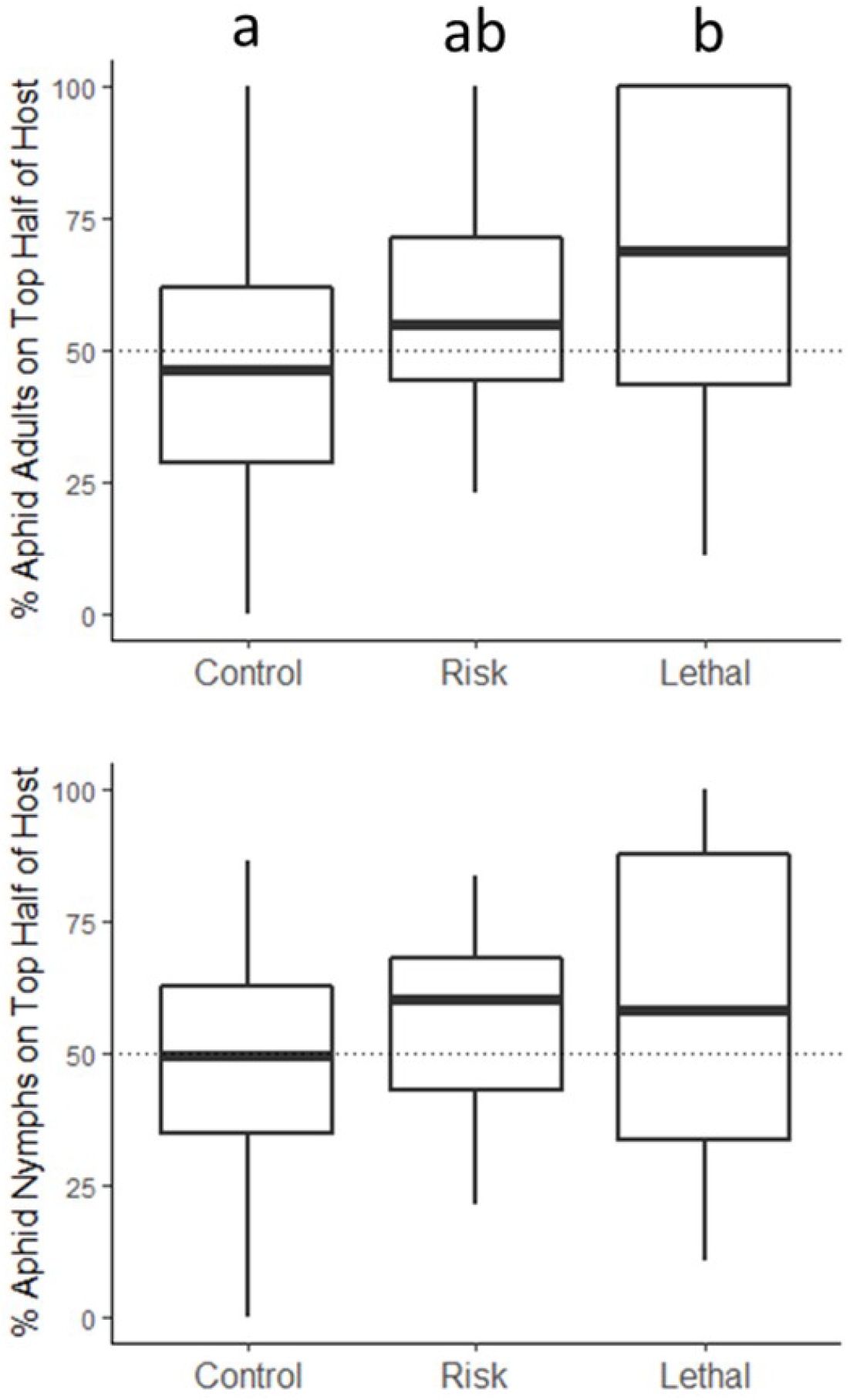
Effects of predator treatments on the feeding location of aphid adults and nymphs on pea hosts. Dotted lines indicate equal numbers of aphids feeding on top vs bottom half of hosts. Letters represent significant differences between groups (Tukey’s HSD, α = .05).

## Discussion

Our study confirms predictions that predation can mediate the transmission dynamics of vector-borne pathogens by affecting aphid abundance and movement. We show the induction of specific aphid behaviors by predators can either promote or interfere with virus transmission. Though lethal predators reduced aphid abundance, the strongest predictor of PEMV prevalence in our model, their effects on aphid movement and feeding behavior appear to have counteracted these reductions, resulting in a net increase in PEMV prevalence when predators were present compared to absent. In contrast, risk-only predators had minimal effects on aphid behaviors but affected PEMV prevalence positively through density-dependent mechanisms. These results lend support to model predictions showing that species interactions affecting vector behavior can contribute more to the rates of vector-borne pathogen spread in ecosystems than those affecting vector abundance (Jeger et al. 2011, Crowder et al. 2019). However, our study shows that ‘risk’ manipulations alone may fail to capture the full range of vector anti-predator responses, which can include responses like compensatory reproduction by herbivores.

Insect herbivores make movement and foraging decisions in response to a ‘landscape of fear’, where tradeoffs between habitat quality and perceived predation risk are weighed (Gaynor et al. 2019). As most plant viruses are dependent on vector dispersal for transmission between hosts (Fereres and Moreno, 2009), responses of herbivores to landscapes of fear are expected to affect pathogen transmission (Finke 2012, Crowder et al. 2019). In our system, lethal predation increased movement of adult aphids, both directly and via reduced aphid abundance, though adult dispersal was not directly linked to increased PEMV. Nymph dispersal, however, may better represent where infectious adult vectors spent time feeding, as nymphs themselves are less likely to leave hosts when threatened by predators (Losey and Denno, 1998). Our results indeed show nymph movement was positively linked with PEMV prevalence, suggesting transmission increased when infectious pea aphids moved to new hosts.

Our model also identified a direct effect of lethal predators on PEMV prevalence, which suggests an important effect on aphids was missing from our *a priori* model, as predators themselves cannot transmit PEMV. Noticing differences in aphid feeding location on host plants between predator treatments in our field study, we recorded feeding location in our greenhouse study. Competitive displacement of aphids to higher locations on individual plants by non-vector herbivores can increase the likelihood of PEMV transmission by causing vectors to feed on more susceptible young tissue (Chisholm et al. 2019). Our results suggest predators may similarly affect virus transmission by displacing aphid vectors upwards on plants (Fig. 3), though such effects could vary in other contexts based on host plant structure and aphid refuge-seeking behavior (Grevstad and Klepetka, 1992, Costamagna and Landis, 2011, Northfield et al. 2012).

Predation risk may induce compensatory responses in insects to defer the reproductive and developmental costs of anti-predator behaviors until risk has decreased (Barribeau et al. 2010, Thaler et al. 2012, Lund 2020). Contrary to our predictions, risk predation increased aphid nymph abundance (Fig. 2), although any long-term negative consequences of compensatory reproduction may not be apparent in our study. It is also possible that disturbance by risk predators induced early dispersal from hosts before aphids reached high densities, at which point reproduction can stall (Agrawal et al. 2004). Moreover, given risk predators’ inability to kill prey, it is possible they did not induce alarm pheromone release by aphid vectors as substantially as lethal predators (Basu et al., 2021). Thus, gross effects of predators reducing vector abundance may be underestimated if non-consumptive effects increase vector abundance.

Risk-manipulated predators performing differently than we predicted highlights the difficulty in establishing how community interactions might affect pathogen transmission *a priori*, as aphid responses to other species can vary based on ecological and environmental context. Aphid’s propensity to disperse or drop from hosts when threatened can be affected by colony density, host plant quality, clonal differences, and previous exposure to risk for example (Kersch-Becker and Thaler, 2015, Muratori et al. 2014, Tamai and Choh, 2019). Moreover, aphid perception of the severity of predation risk is dependent on predator foraging strategies, recognition of risk signals, and shared evolutionary history (Preisser et al. 2007, Sih et al. 2010). Given the potential for interactions and feedback among prey responses to predators (Sheriff et al. 2020), inclusion of multiple vector responses in analyses remains critical to identifying mechanisms behind observed changes in transmission.

Our study provides an experimental and statistical framework for the examination of a broader range of predator effects on vectors and pathogen transmission. Variation in predator communities or density would likely affect the relative magnitude of predators’ effects on vector abundance, development, and behaviors (Finke, 2012). Diverse predator communities can act synergistically or antagonistically in suppressing herbivore populations (Losey and Denno, 1998, Snyder et al. 2001, 2008), though the effects of predator diversity on vector behavior and virus transmission have been poorly investigated. Additionally, pathogen characteristics determine the importance of specific vector responses to overall rates of transmission; vector behaviors that accelerate the transmission of certain pathogens (e.g. rapid probing and dispersal) may stall the transmission of others (Long and Finke, 2015, Mauck et al. 2018, Crowder et al. 2019). Our results and other accumulating evidence suggest vector abundance alone may not be a suitable indicator for disease risk in natural systems, and future studies manipulating the characteristics of pathosystems will help to identify the mechanisms through which trophic interactions can affect vector-borne pathogen transmission.

## Acknowledgements

We thank the undergraduates who assisted with experiments and T. Cutter and M. Asche for illustrations. This work was supported by USDA NIFA Award 2019-67011-29602.

## References

Agrawal, A. A. et al. 2004. Intraspecific variation in the strength of density dependence in aphid populations. – Ecol. Entomol. 29: 521–526.

Barribeau, S. M. et al. 2010. Aphid reproductive investment in response to mortality risks. – BMC Evol. Biol. 10: 251

Basu, S. et al. 2021. Insect alarm pheromones in response to predators: Ecological trade-offs and molecular mechanisms. – Insect Biochem. Mol. Biol. 128: 103514.

Bates, D. et al. 2015. Fitting linear mixed-effects models using lme4. – J. Stat. Softw. 67

Bowers, W. S. et al. 1972. Aphid alarm pheromone: Isolation, identification, synthesis. – Science 177: 1121–1122.

Chisholm, P. J. et al. 2019. Plant-mediated interactions between a vector and a non-vector herbivore promote the spread of a plant virus. – Proc. R. Soc. B Biol. Sci. 286: 20191383.

Clark, R. E. et al. 2019. Tri-trophic interactions mediate the spread of a vector-borne plant pathogen. – Ecology 100: e02879

Costamagna, A. C. and Landis, D. A. 2011. Lack of strong refuges allows top-down control of soybean aphid by generalist natural enemies. – Biol. Control 57: 184–192.

Crowder, D. W. et al. 2019. Species interactions affect the spread of vector-borne plant pathogens independent of transmission mode. - Ecology 100: 1–10.Fan, L. et al. 2017. Adaptation of Defensive Strategies by the Pea Aphid Mediates Predation Risk from the Predatory Lady Beetle. J. Chem. Ecol44: 40–50

Fereres, A., and A. Moreno. 2009. Behavioural aspects influencing plant virus transmission by homopteran insects 141:158–168.

Finke, D. L. 2012. Contrasting the consumptive and non-consumptive cascading effects of natural enemies on vector-borne pathogens. – Entomol. Exp. Appl. 144: 45–55.

Fox J, Weisberg S. 2019. An R Companion to Applied Regression, Third edition. Sage, Thousand Oaks CA.

Gaynor, K. M., J. S. Brown, A. D. Middleton, M. E. Power, and J. S. Brashares. 2019. Landscapes of FearLJ: Spatial Patterns of Risk Perception and Response. Trends in Ecology & Evolution 34:1–14.

Grevstad, F. S. et al. 2014. The Influence of Plant Architecture on the Foraging Efficiencies of a Suite of Ladybird Beetles Feeding on Aphids. Oecologia 92: 399–404.

Hodge, S. et al. 2011. Parasitoids aid dispersal of a nonpersistently transmitted plant virus by disturbing the aphid vector. – Agric. For. Entomol. 13: 83–88.

Jeger, M. J. et al. 2011. Interactions in a host plant-virus – vector – parasitoid systemLJ: Modelling the consequences for virus transmission and disease dynamics. – Virus Res. 159: 183–193.

Jones, E. I. and Dornhaus, A. 2011. Predation risk makes bees reject rewarding flowers and reduce foraging activity. – Behav. Ecol. Sociobiol. 65: 1505–1511.

Kersch-Becker, M. F. and Thaler, J. S. 2015. Plant resistance reduces the strength of consumptive and non-consumptive effects of predators on aphids. – J. Anim. Ecol. 84: 1222– 1232.

Lefcheck, J. S. 2016. piecewiseSEM: Piecewise structural equation modelling in r for ecology, evolution, and systematics. – Methods Ecol. Evol. 7: 573–579.

Lenth, R. 2020. emmeans: estimated marginal means, aka least-squares means. – R package ver. 1.5.0.

Long, E. Y. and Finke, D. L. 2015. Predators indirectly reduce the prevalence of an insect vectored plant pathogen independent of predator diversity. – Oecologia 177: 1067–1074.

Losey, J. E. and Denno, R. F. 1998. The escape response of pea aphids to foliar-foraging predators: Factors affecting dropping behaviour. – Ecol. Entomol. 23: 53–61.

Lund, M. et al. 2020. Predation threat modifies Pieris rapae performance and response to host plant quality. – Oecologia 193: 389–401

Mauck, K. E. et al. 2018. Evolutionary Determinants of Host and Vector Manipulation by Plant Viruses. – In: Advances in Virus Research. Elsevier Inc, pp. 189–250

Moore, S. M. et al. 2010. Predators indirectly control vector-borne disease: linking predator-prey and host-pathogen models. – J. R. Soc. Interface 7: 161–176.

Muratori, F. B. et al. 2014. Clonal variation in aggregation and defensive behavior in pea aphids. – Behav. Ecol. 25: 901–908.

Northfield, T. D. et al. 2012. A simple plant mutation abets a predator-diversity cascade. – Ecology 93: 411–420.

Preisser, E. L. et al. 2005. Scared to Death? The Effects of Intimidation and Consumption In Predator-Prey Interactions. – Ecology 86: 501–509.

Preisser, L. et al. 2007. Predator Hunting Mode and Habitat Domain Alter Nonconsumptive Effects in Predator-Prey Interactions. – Ecology 88: 2744–2751.

Sheriff, M. et al. 2020. Non-consumptive predator effects on prey population size: A dearth of evidence. – J. Anim. Ecol. 89: 1302–1316.

Sih, A. et al. 2010. Predator – prey naïveté, antipredator behavior, and the ecology of predator invasions. Oikos 119: 610–621.

Smyrnioudis, I. N. et al. 2001. The effect of natural enemies on the spread of barley yellow dwarf virus (BYDV) by Rhopalosiphum padi (Hemiptera: Aphididae). – Bull. Entomol. Res. 91: 301–306.

Snyder, W. E. and Ives, A. R. 2001. Generalist predators disrupt biological control by a specialist parasitoid. – Ecology 82: 705–716.

Snyder, G. B. et al. 2008. Predator biodiversity strengthens aphid suppression across single- and multiple-species prey communities. – Biol. Control 44: 52–60.

Tamai, K. and Choh, Y. 2019. Previous exposures to cues from conspecifics and ladybird beetles prime antipredator responses in pea aphids Acyrthosiphon pisum (Hemiptera: Aphididae). – Appl. Entomol. Zool. 54: 277–283.

Thaler, J. S. et al. 2012. Compensatory mechanisms for ameliorating the fundamental trade-off between predator avoidance and foraging. – Proc. Natl. Acad. Sci. U. S. A. 109: 12075– 12080.

Tholt, G. et al. 2018. Could vectors’ fear of predators reduce the spread of plant diseases? – Sci. Rep. 8: 8705.

