## Supplemental Figures and Tables for "Predators affect a plant virus through direct and trait-mediated indirect effects on vectors"

Figure S1. Field experiment layout with quadrant sub-sampling. Aphids were initially confined to the center plant (green square). After the experiment, all plants (orange, blue, and green squares) were visually evaluated for PEMV symptoms. Three plants from within each quadrant (plants in orange, quadrant shown by red box) were destructively sampled and tested for PEMV titer using rt-PCR.

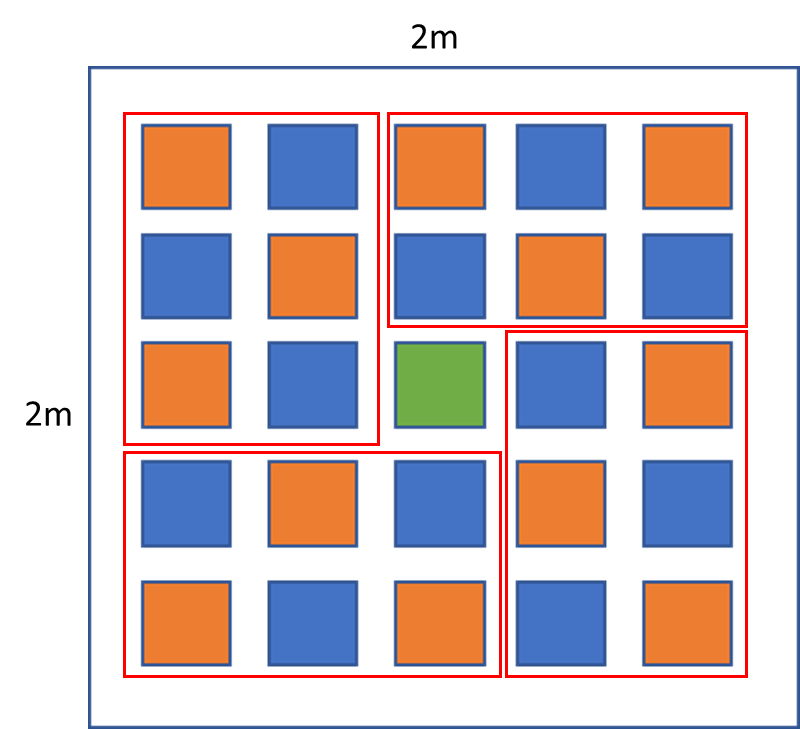

Figure S2. Electrophoresis Gels based on amplification products of PEMV-1 coat protein. Labels corresponds to treatment replicates and quadrants of experimental mesocosms within which 3 plants were subsampled and pooled for analysis, with identical positive(+) and negative(–) controls run on each gel. ImageJ was used to quantify band intensity, and quadrant titer was evaluated relative to positive control.

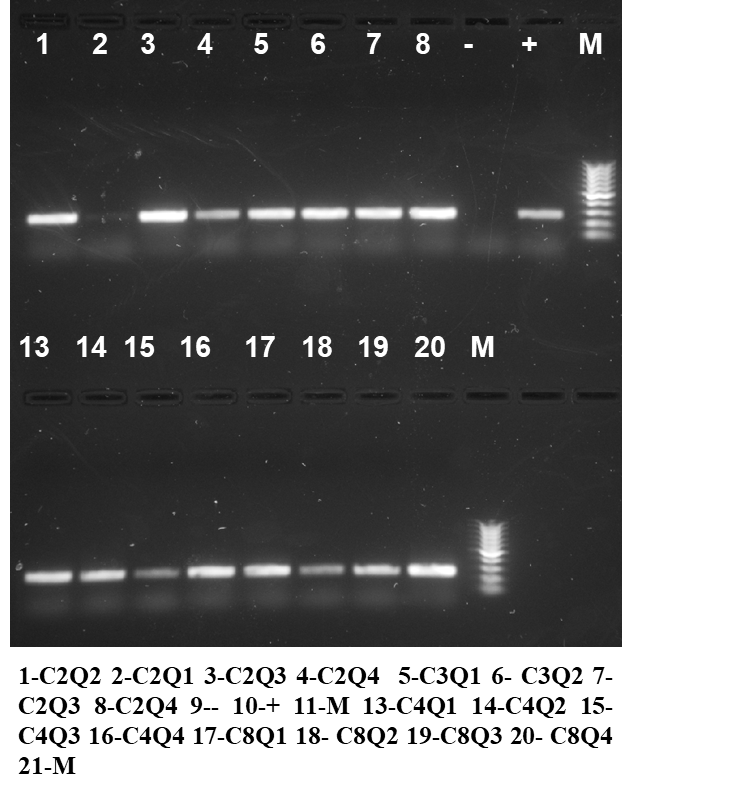

Figure S3. Aphid adult and nymph abundance (a,b) and average distance from initial host in inches (c,d) over time in response to predation treatments. Error bars indicate standard error of the mean.

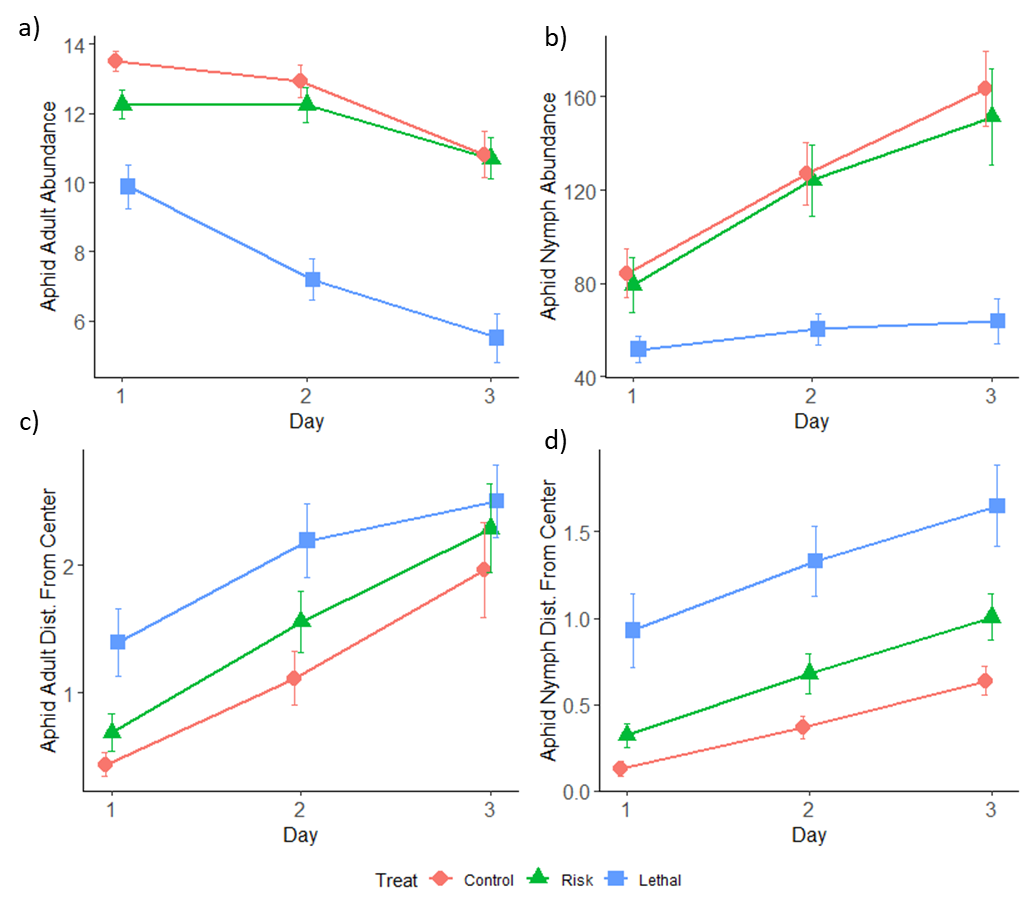

Table S1. Standardized path coefficients, significance tests, and tests of directed separation from accepted path model. Standardized coefficients (βstd) indicate relative magnitude and direction of effect. Predictors and coefficients under heading of ‘Tests of directed separation' indicate dropped interactions that did not significantly contribute to path model.

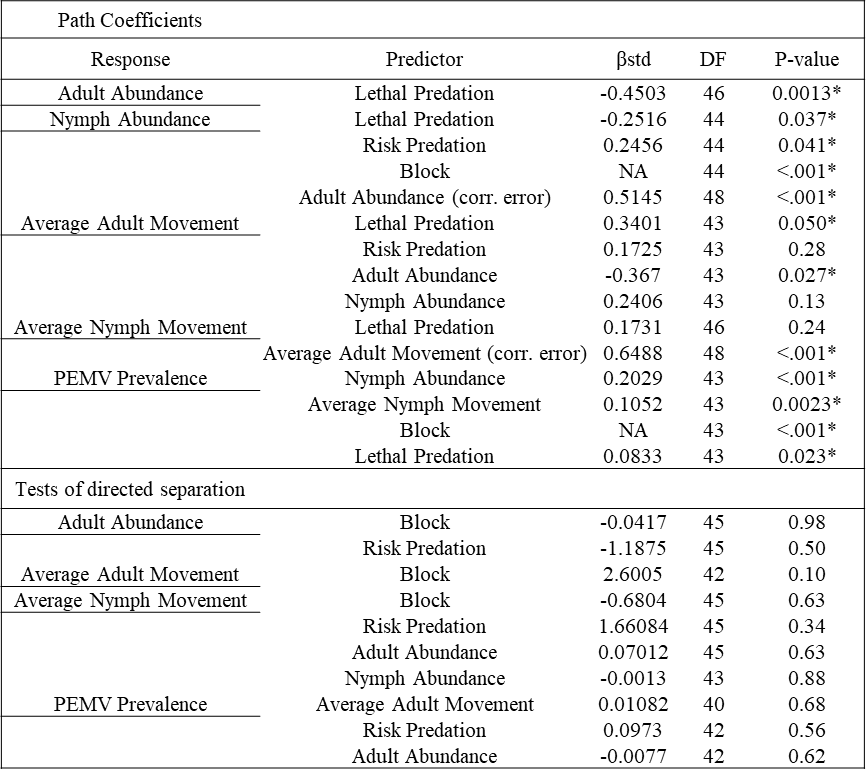

Table S2. Model Specifications and significance tests for GLMMs on aphid responses to predation over time.

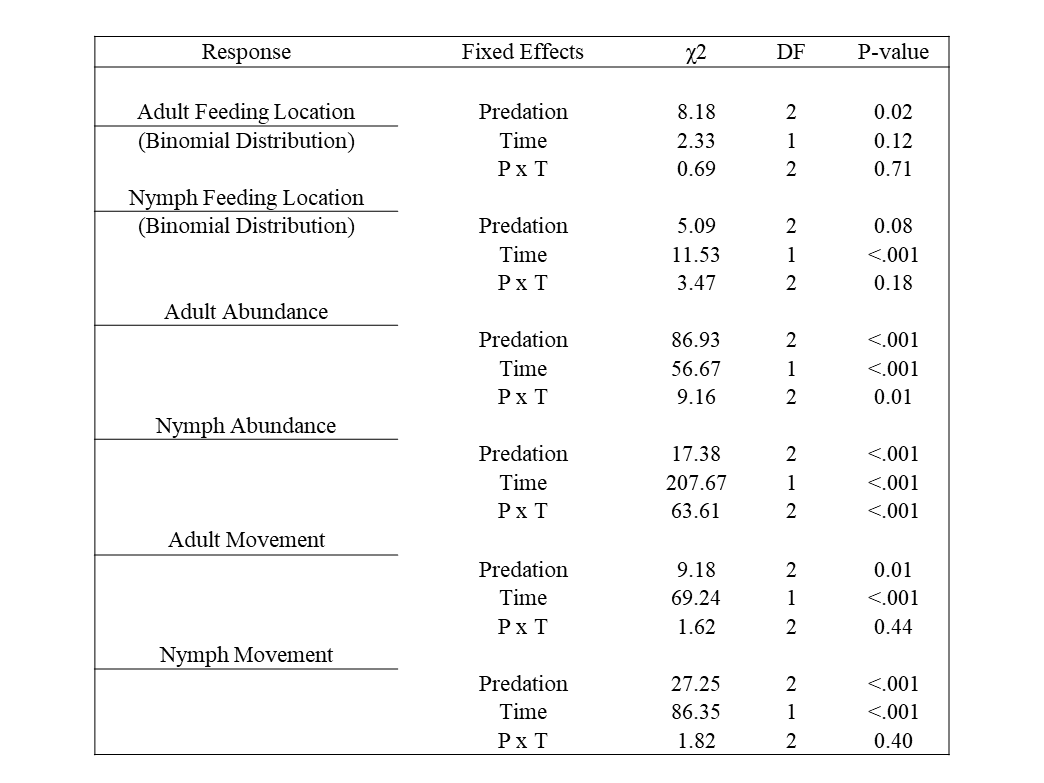
